# Koisio Technology-Modulated Cell Culture Media can Significantly Increase the Antioxidant Capacity of Mouse Fibroblasts

**DOI:** 10.1101/2022.03.03.482792

**Authors:** Mingchao Zhang, Yinghui Men, Qi Zhu, Weihai Ying

**Author notes:** These two authors contributed equally to this work (Mingchao Zhang, Yinghui Men). Corresponding author, Weihai Ying, Ph.D. Professor, School of Biomedical Engineering and Med-X Research Institute, Shanghai Jiao Tong University, 1954 Huashan Road, Shanghai, 200030, P.R. China.

## Abstract

Oxidative stress is not only a critical common pathological factor of numerous diseases, but also an important factor in the aging process. It is highly valuable to discover novel and economic substances for enhancing the antioxidant capacity of human body. In this study we used a cell culture model to determine the antioxidant capacity of Koisio technology-modulated solutions without additions of any external substances. Our study has obtained the following findings: First, the cells cultured in Koisio technology-modulated cell culture media (KM) showed significantly greater capacity to decrease H_2_O_2_-induced oxidative stress, compared with the cells cultured in regular cell culture media; second, the cells cultured in KM showed significantly greater capacity to resist H_2_O_2_-induced decreases in cell number, compared with the cells cultured in regular cell culture media; and third, the H_2_O_2_ prepared in Koisio technology-modulated MEM produced significantly lower levels of decreases in cell number, compared with the H_2_O_2_ prepared in MEM. Collectively, our study has indicated that Koisio technology can increase the antioxidant capacity of both solutions and cells without additions of any external substances. Based on our previous findings, we proposed that Koisio technology-modulated solutions increase cellular antioxidant capacity at least partially by increasing intracellular ATP levels.

## Introduction

Oxidative stress is a critical common pathological factor of numerous diseases, including acute ischemic stroke (AIS) ^1,2^, cardiovascular diseases ^3^, lung cancer ^4^, Alzheimer’s disease ^5^, and Parkinson’s disease ^6^, but also an important factor in the aging process. While our understanding on the mechanisms underlying oxidative damage has been greatly improved through numerous studies, many critical mechanisms of oxidative damage have remained unclear.

It is highly valuable to discover novel and economic substances for enhancing the antioxidant capacity of human body. Since water drinking is required for human survival, it would be of great value if we can produce certain type of water with antioxidant capacity without addition of external substances. There have been some types of drinking water that has shown antioxidant capacity. However, most of these types of drinking water requires additions of external substances. These types of water have significant limitations, including relatively low stability of the active components in the water or limited resource of the water.

Koisio technology is a novel technology that modulates water solely by physical approaches without additions of any external substances. A previous study has reported that the ceramics produced by Koisio technology could produce water with significantly increased permeability through aquaporins ^7^. Our latest study has shown the capacity of Koisio technology-modulated media to markedly increase intracellular ATP levels ^8^. Our study has also found that Koisio technology-modulated media can promote growth of lung fibroblasts ^9^. However, it has been unclear if Koisio technology may affect the antioxidant capacity of solutions and cells. In this study we used mouse fibroblast L929 cells as a cellular model to determine if Koisio technology may affect antioxidant capacity of solutions and cells. Our study has indicated that Koisio technology can increase the antioxidant capacity of both solutions and cells without additions of any external substances.

## Materials and Methods

### Regents

#### Cell Cultures

Mouse lung fibroblast (L929) cell lines ^10, 11^ were purchased from Chinese Academy of Sciences Cell Bank. The cells were grown in DMEM (SH30243.01, Hyclone) supplemented with 10% FBS (100-500, GEMINI), 100 U/mL penicillin, and 0.1mg/mL streptomycin (15140122, Gibco) at 37 °C in humidified 5% CO_2_ atmosphere.

### Modulations of cell culture media by Koisio technology

Modulations of cell culture media were conducted by Koisio technology as reported previously ^7^.

### Determinations of intracellular ROS levels

For Dichlorofluorescin (DCF) assay, 2,7-Dichlorofluorescin diacetate (DCFH-DA, Beyotime, China), a ROS-specific fluorescent probe, was used to measure total intracellular ROS levels. Cells were incubated with 20 μM DCFH-DA dissolved in DMEM without fetal bovine serum (FBS) for 20 min at 37°C in an incubator. After 3X washes with PBS, the cells were observed under a fluorescence microscope (Leica, Wetzlar, Germany) or analyzed by flow cytometry (FACS Aria; Becton Dickinson, Heidelberg, Germany) to detect the mean fluorescence intensity (MFI) with an excitation wavelength of 488 nm and emission wavelength of 525 nm.

### Intracellular lactate dehydrogenase (LDH) assays

Cell survival was quantified by measuring the intracellular LDH activity of the cells. Intracellular LDH activity was determined by measuring the LDH activity of cell lysates. Cell lysates were prepared through the following procedures: Cells were lysed for 15 min in lysing buffer containing 0.04% Triton X-100, 2 mM HEPES and 0.01% bovine serum albumin (pH 7.5). For LDH activity assays, 50 μL cell culture medium or 50 μL cell lysates was mixed with 150 μL 500 mM potassium phosphate buffer (pH 7.5) containing 0.34 mM NADH and 2.5 mM sodium pyruvate. The A_340nm_ changes were monitored over 90 sec. The percentage of cell survival was calculated by normalizing the LDH values of samples to LDH activity measured in the lysates of control (wash only) culture wells.

### Statistical analyses

All data are presented as mean + SEM. Data were assessed by one-way ANOVA, followed by Student - Newman - Keuls post hoc test. *p* values less than 0.05 were considered statistically significant.

## Results

### The L929 cells cultured in the Koisio technology-based DMEM cell culture media (KM) showed significantly greater capacity to decrease ROS levels in the H_2_O_2_-treated L929 cells, compared with the cells cultured in regular cell culture media

After L929 cells were cultured in either regular cell culture media or KM for 6 - 15 generations, the cells were treated with H_2_O_2_. FACS-based assay using DCFH as ROS probe showed that H_2_O_2_ led to significant increases in DCF signals in both types of cell cultures (Fig. 1A and Fig. 1B). The intracellular ROS level in the H_2_O_2_-treated cells cultured in KM was significantly lower than that in the H_2_O_2_-treated cells cultured in regular cell culture media (Fig. 1A and Fig. 1B).

**Fig. 1.**
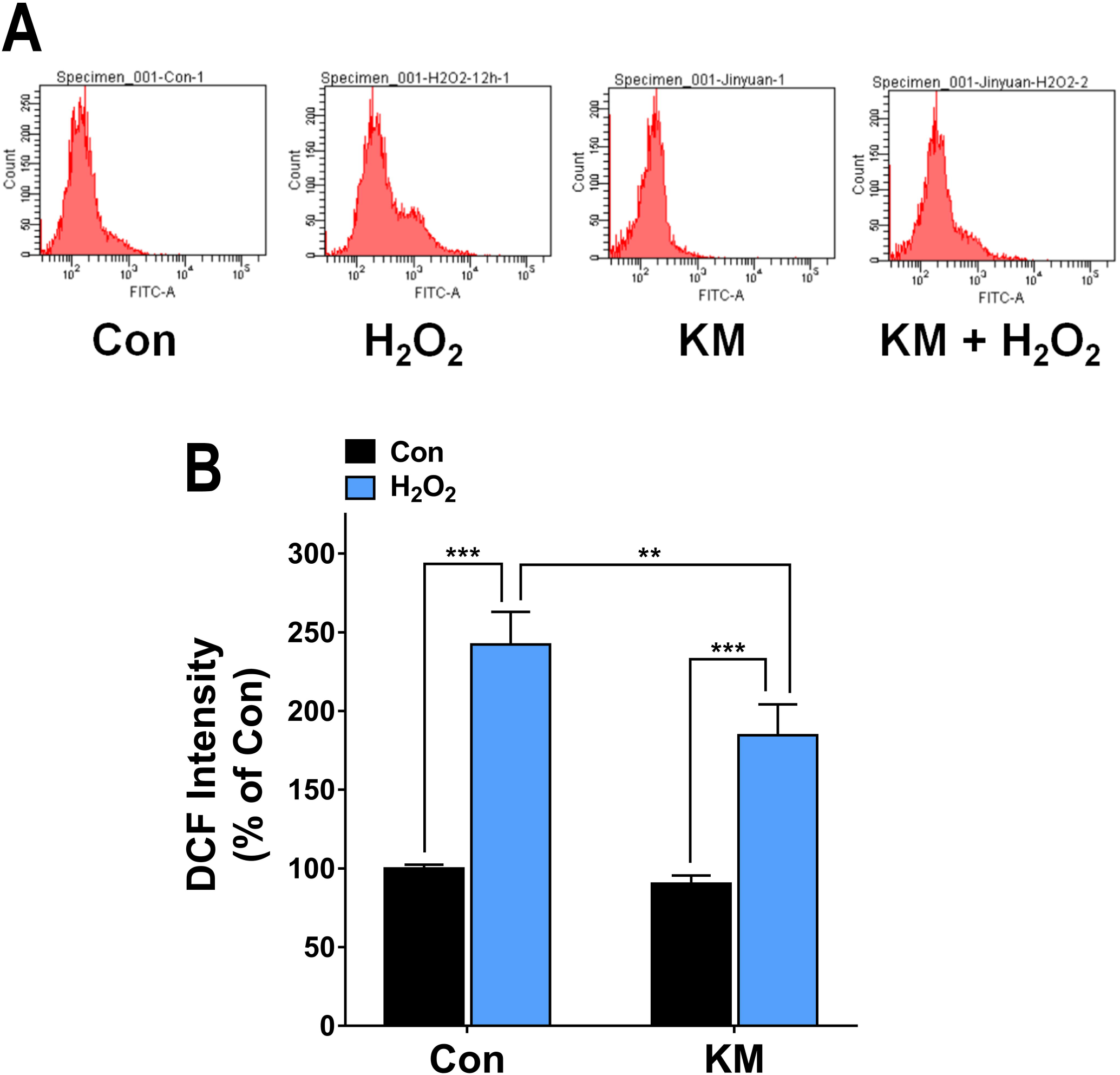
The L929 cells cultured in the Koisio technology-based DMEM cell culture media (KM) showed significantly greater capacity to decrease ROS levels in the H_2_O_2_-treated mouse fibroblasts, compared with the cells cultured in regular cell culture media. After L929 cells were cultured in either regular cell culture media or KM for 6 - 15 generations, the cells were exposed to the H_2_O_2_ prepared in MEM for 45 min. After the treatment, the cells were cultured in regular cell culture media or KM for 18 – 24 hrs. Subsequently FACS-based assay using DCFH as ROS probe was conducted. The FACS-based assay showed that the intracellular ROS level in the cells cultured in KM was significantly lower than that in the cells cultured in regular cell culture media. **, *P* < 0.01; ***, *P*< 0.001. N = 8 - 9. The data were collected from three independent experiments.

### The L929 cells cultured in KM showed significantly greater capacity to resist H_2_O_2_-induced decrease in cell number, compared with the cells cultured in regular cell culture media

After L929 cells were cultured in either regular cell culture media or KM for 6 −15 generations, the cells were treated with H_2_O_2_. Intracellular LDH assay showed that H_2_O_2_ led to significant decreases in the intracellular LDH activity of both types of cells (Fig. 2). The intracellular LDH activity of the H_2_O_2_-treated cells cultured in KM was significantly higher than that of the H_2_O_2_-treated cells cultured in regular cell culture media (Fig. 2).

**Fig. 2.**
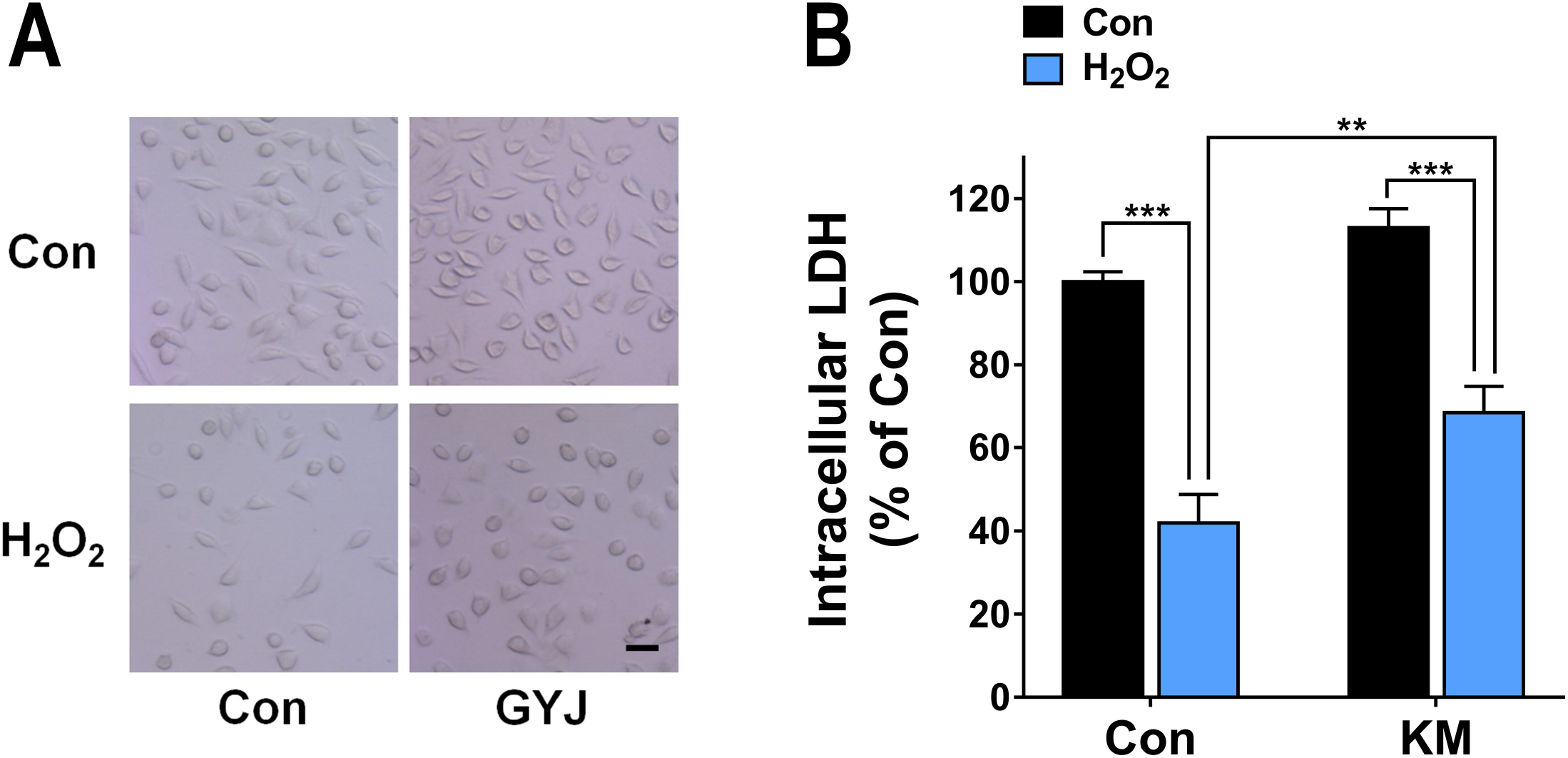
The L929 cells cultured in KM showed significantly greater capacity to resist H_2_O_2_-induced decrease in cell number, compared with the cells cultured in regular cell culture media. After L929 cells were cultured in regular cell culture media or KM for 6 - 15 generations, the cells were exposed to the H_2_O_2_ prepared in MEM for 45 min. After the treatment, the cells were cultured in regular cell culture media or KM for 18 – 24 hrs. Subsequently intracellular LDH assay was conducted. Intracellular LDH assay showed that the L929 cells cultured in KM showed significantly greater capacity to resist H_2_O_2_-induced decrease in cell number, compared with the cells cultured in regular cell culture media. ***, *P* < 0.001. N = 9. The data were collected from three independent experiments.

### The H_2_O_2_ prepared in Koisio technology-modulated MEM (K-MEM) produced decreases in cell number to a significant smaller degree compared with the H_2_O_2_ prepared in MEM

After L929 cells were cultured in regular cell culture media for 6 −15 generations, the cells were treated with the H_2_O_2_ prepared in MEM or the H_2_O_2_ prepared in K-MEM. Intracellular LDH assay showed that compared with the cells treated with the H_2_O_2_ prepared in MEM, there was a significantly higher level of intracellular LDH activity in the cells treated with the H_2_O_2_ prepared in Koisio technology-modulated MEM (Fig. 3).

**Fig. 3.**
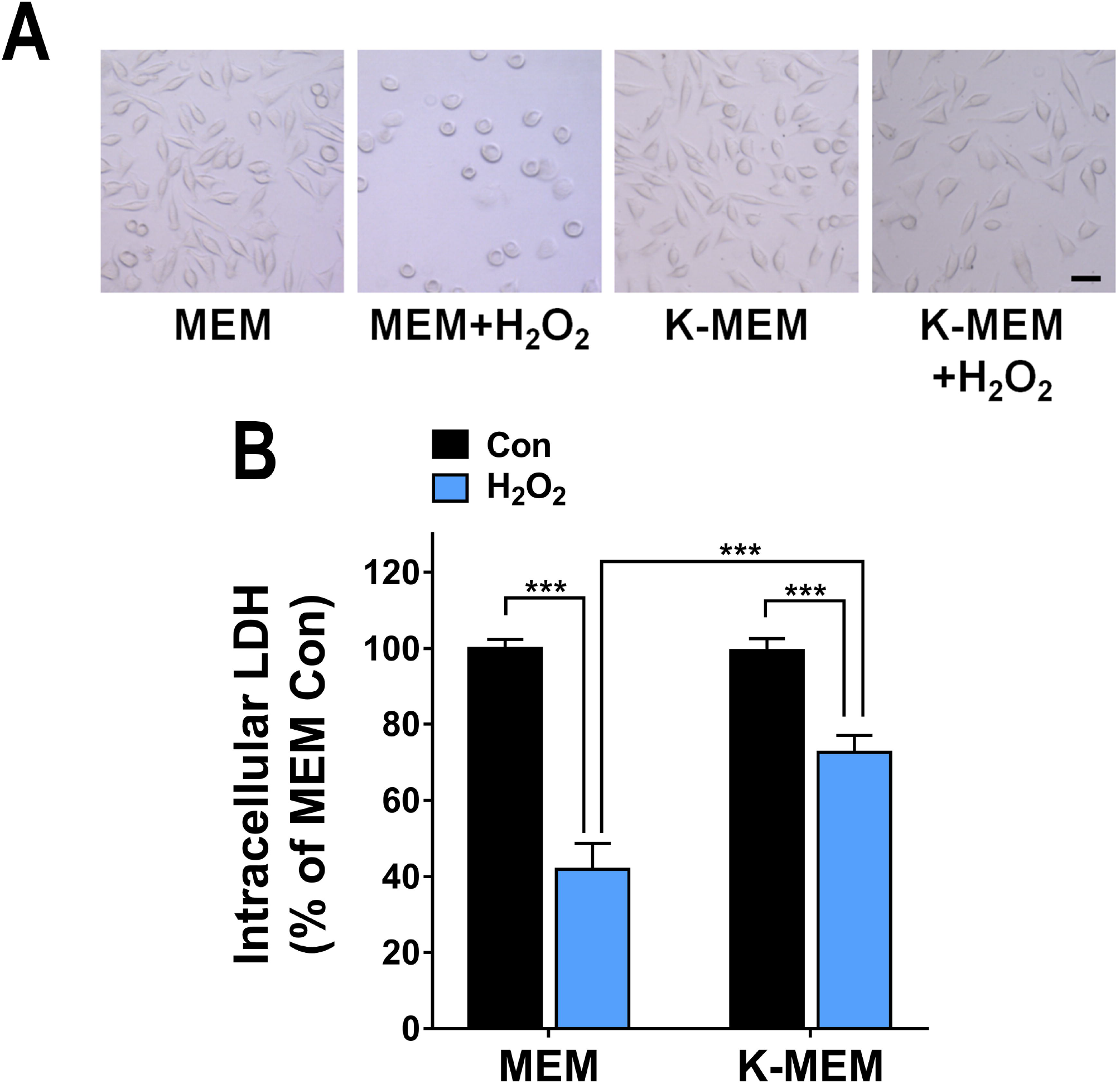
The H_2_O_2_ prepared in Koisio technology-modulated MEM (K-MEM) produced decreases in cell number to a significant smaller degree compared with the H_2_O_2_ prepared in MEM. After L929 cells were cultured in regular cell culture media for 6 - 15 generations, the cells were exposed to the H_2_O_2_ prepared in MEM or the H_2_O_2_ prepared in K-MEM for 45 min. After the treatment, the cells were cultured in regular cell culture media for 18 – 24 hrs. Subsequently intracellular LDH assay was conducted. Intracellular LDH assay showed that compared with the cells treated with the H_2_O_2_ prepared in MEM, there was a significantly higher level of intracellular LDH activity in the cells treated with the H_2_O_2_ prepared in K-MEM. ***, *P* < 0.001. N = 9. The data were collected from three independent experiments.

## Discussion

The major findings of our study include: First, the cells cultured in KM showed significantly greater capacity to decrease H_2_O_2_-induced oxidative stress, compared with the cells cultured in regular cell culture media; second, the cells cultured in KM showed significantly greater capacity to resist H_2_O_2_-induced decreases in cell number, compared with the cells cultured in regular cell culture media; and third, the H_2_O_2_ prepared in Koisio technology-modulated MEM produced a significantly lower level of decreases in cell number, compared with the H_2_O_2_ prepared in MEM.

Two lines of evidence has suggested that Koisio technology can lead to significant increases in the antioxidant capacity of cells without additions of any external substances: First, the cells cultured in KM showed significantly greater capacity to decrease H_2_O_2_-induced oxidative stress; and second, the cells cultured in KM showed significantly greater capacity to resist H_2_O_2_-induced decreases in cell number. One line of evidence has suggested that Koisio technology can lead to significant increases in the antioxidant capacity of solutions: The H_2_O_2_ prepared in Koisio technology-modulated MEM produced a significantly lower level of decreases in cell number.

It is important to elucidate the mechanisms underlying the effects of KM on the antioxidant capacity of cells. Since our previous study has found that KM can produce a remarkable increase in the intracellular ATP levels ^8^, KM may produce the increased antioxidant capacity by the following mechanisms: Increased intracellular levels can lead to inhibition of glycolysis through its inhibitory effects on the rate-limiting glycolytic enzymes ^12^. The glycolytic inhibition can lead to increased flow of glucose-6-phosphate into pentose phosphate pathway, resulting in increased production of NADPH – a key antioxidant molecule that is required for regeneration of GSH ^12^.

We found that the H_2_O_2_ prepared in the Koisio technology-modulated MEM led to a significantly lower degree of decreased cell number, compared with the H_2_O_2_ prepared in the regular MEM. This study has suggested that the Koisio technology-modulated MEM has direct antioxidant capacity on H_2_O_2_. It is warranted to further investigate the mechanisms underlying the antioxidant capacity of Koisio technology-modulated solutions.

## Notes

### Competing Interest Statement

The authors have declared no competing interest.

